# The Dorsolateral Prefrontal Cortex Presents Structural Variations Associated with Empathic Capacity in Psychotherapists

**DOI:** 10.1101/2021.01.02.425096

**Authors:** Marcos E. Domínguez-Arriola, Víctor E. Olalde-Mathieu, Eduardo A. Garza-Villareal, Fernando A. Barrios

## Abstract

Empathic capacity has been shown to be correlated with brain structural variations, such as cortical thickness. Since psychotherapists have a constant demand to modulate their empathic response, in this study we compared cortical thickness between a group of psychotherapists and a control group at prefrontal and cingulate brain regions, and investigated how this is correlated with their empathic skills. Eighteen psychotherapists and eighteen healthy controls underwent 3-Tesla MRI scanning and completed empathy-related psychometric assessments. Cortical thickness (CT) measures were estimated for each participant. We evaluated how these measures differed between groups, and if they were associated with individual empathy-related scores in a series of regions of interest. Our analysis shows that psychotherapists display a significantly greater CT at a region in the left dorsolateral prefrontal cortex (dlPFC; p < 0.05, FDR corrected). Moreover, psychotherapists’ CT in this region is correlated with the tendency to feel empathically concerned for others (p < 0.01, FDR corrected). This finding is relevant because the dlPFC region participates strongly in the cognitive components of the empathic response, such as emotion regulation and perspective-taking processes. Thus, our findings support the idea that empathic capacity is reflected by brain structural variations while also studying for the first time a sample of subjects for whom empathic responding is crucial in their profession.

## Introduction

Recently, there has been increasing interest in how the adult brain’s plasticity is susceptible to individual differences in social processing capacities. In this context, the effect of empathy has received significant attention. Empathy is defined as an umbrella term that encompasses all processes through which an organism—generally a mammal—can understand others’ affective states through the activation of their own representations of such states (Bernhardt & Singer 2012; De Waal & Preston 2017; Preston & De Waal 2002). Such processes engage a range of brain regions including the amygdala, the cingulate cortex and some prefrontal regions (De Waal & Preston 2017). What is more, some of these regions display structural variations as a function of different empathy measures (Banissy et al. 2012; Eres, Decety & Louis 2015; Uribe, Puig-Davi, Abos et al. 2019).

Empathic processes are generally classified into two categories. Affective empathy phenomena are bottom-up processes which depend on the coupling of perception and action, and engage brain regions associated with first-hand affective experience and sensori-motor activity—like insula, amygdala, somatosensory and motor cortices and ventromedial prefrontal cortex (vmPFC). Whereas cognitive empathy phenomena are top-down, executive processes engaging brain structures that participate in mentalizing and working memory—such as dorsolateral prefrontal cortex (dlPFC), vmPFC, and posterior cingulate cortex (pCC). (Bernhardt & Singer 2012; De Waal & Preston 2017; Engen & Singer 2013)

Nonetheless, these distinct empathic processes do not happen independently. Rather, they remain functionally integrated and a real-life empathic response will engage both affective and cognitive empathic phenomena (De Waal & Preston 2017; Zaki et al. 2009). Accordingly, Decety (2011) and his colleagues have proposed that an empathic response consists of three successive components: affective arousal, emotional awareness, and emotion regulation. While the first component represents the bottom-up (“affective”) aspect of the empathic response, its emotion regulation component characterizes its most top-down (“cognitive”) component and permits effective engagement in helping behavior (Decety 2011; Elliot et al. 2011; Lamm et al. 2007a).

Emotion regulation refers to the modulation of ongoing affective states through conscious or unconscious processes. (Etkin, Büchel & Gross 2015; Ochsner & Gross 2005). A distinction between processes of behavioral and cognitive emotion regulation is commonly made (Ochsner & Gross 2005). While the former refer to inhibition of expressive behavior (i.e. expressive suppression), the latter point to top-down mechanisms aimed to modulate the intensity of the emotional experience, or change the emotion itself (e.g. attentional deployment away from, or cognitive reappraisal of, the eliciting stimuli). (Etkin, Büchsel & Gross 2015; Gross 2002; Ochsner & Gross 2005)

Psychotherapists might suppress the expression of their emotions during a psychotherapy session as a first resource so as not to display their full emotional response to the patient (for instance, when the therapist becomes angry) (Prikhidko & Swank 2018). However, this type of inhibition tends to impair some aspects of cognition such as memory about the emotion-eliciting stimuli, and also has negative effects on the ongoing social interaction (Gross, 2002). As a result, expressive suppression is not a desirable emotion regulation strategy for psychotherapists (Pletzer, Sanchez & Scheibe 2015; Prikhidko & Swank 2018).

Cognitive emotion regulation mechanisms do not seem to have such detrimental effects on cognition and social interaction, probably because the intensity of the emotional experience itself is directly modulated (Gross, 2002). This makes cognitive emotion regulation more suitable for psychotherapists’ empathic responding (Pletzer, Sanchez & Scheibe 2015; Prikhidko & Swank 2018). There is consistent evidence that this kind of regulation is mediated by prefrontal dorsolateral regions (Etkin, Büchel & Gross 2015; Ochsner & Gross 2005), especially in the left hemisphere (Ochsner et al. 2004) and involving to a lesser extent ventrolateral prefrontal and parietal cortices (Etkin, Büchel & Gross 2015).

Along these lines, the empathic response (especially its cognitive emotion regulation component) has an important role in psychotherapy (Elliot et al. 2011). When psychotherapists effectively modulate their empathic responding and reflect an emotional response that is congruent with the one of the patient, it is more likely that the patient feels understood and cared about, and even that they become more aware of their own affective state (Pletzer, Sanchez & Scheibe 2015). The relevance of this is to such an extent that the level of empathy during the psychotherapy session, as perceived by the patient, has been found to be a robust predictor of therapeutic success, independently of the theoretical system (Elliot et al. 2011; MacFarlane et al. 2017). This finding suggests that there might be a pressure for psychotherapists to be more empathic through continuously modulating their empathic response. The hypothesis that psychotherapists have an increased empathic capacity has been tested by some research groups, finding evidence that, even though psychotherapists show similar levels of bottom-up emotional reactivity as nontherapists (Hassentab et al. 2007; Pletzer et al. 2015), they display a significantly higher cognitive empathy capacity as measured through behavioral tasks and psychometric scales (Hassentab et al. 2007; Olalde-Mathieu et al. 2020).

A further reason for the need of psychotherapists to be especially empathic is that they tend to experience strong emotions of anger, guilt and anxiety as a result of their professional practice. Due to these intense emotions psychotherapists are in constant risk of vicarious traumatization and, consequently, professional impairment. Thus, the constant monitoring of their own affective state, and the use of strategies to regulate it is even a matter of ethical responsibility (Prikhidko & Swank 2018). This means that psychotherapists are required to reliably modulate their emotional response even after work, whether this means down-regulating intense emotions or up-regulating desirable emotional states (Prikhidko & Swank 2018). Accordingly, some studies have found that psychotherapists are better at regulating their emotions than non-therapists (Pletzer et al. 2015), and that they tend to draw less often upon suppression as a strategy to modulate their affective state (Olalde-Mathieu et al. 2020).

That said, however, the impact that this might have in terms of brain plasticity in psychotherapists has not yet been investigated. The assumption that the healthy adult human brain is susceptible to local plasticity through repeated exposure to certain kinds of stimuli and environmental demands is strongly supported by empirical evidence (Banks, Sreenivasan, Weintraub et al. 2016; Bermudez & Zatorre 2005; Delon-Martin, Plailly, Fonlupt et al. 2013; Maguire, Gadian, Johnsrude et al. 2000). These variations are detectable through structural MRI analysis techniques such as voxel-based morphometry (Ashburner & Friston 2000) and cortical thickness analysis (Lerch & Evans 2005; Lerch, Van Der Kouwe, Raznahan et al. 2017).

Such methods have yielded evidence that certain brain structures tend to structurally vary as a function of various empathic measures. While affective empathy measures tend to correlate with structural variation in bilateral insula (Banissy et al. 2012; Eres, Decety & Louis 2015), left anterior cingulate cortex (ACC), left precuneus, and left somatosensory cortex (Banissy et al. 2012), cognitive empathy measures correlate with structural variations in structures that comprise the dlPFC (Banissy et al. 2012; Uribe, Puig-Davi, Abos et al. 2019), left cingulate cortex (CC) (Banissy et al. 2012; Eres, Decety & Louis 2015; Uribe, Puig-Davi, Abos et al. 2019), and dorsomedial prefrontal cortex (dmPFC) (Eres, Decety & Louis 2015; Uribe, Puig-Davi, Abos et al. 2019).

Although the brain structural correlates of emotion regulation skills in healthy adults have been scarcely studied, these have shown to be somewhat distributed throughout the brain. Previous studies have shown gray matter density variations associated to emotion regulation-related measures in ACC, PCC, dlPFC, dmPFC, supplementary motor area, amygdala, precuneus, superior temporal gyrus (STG), and anterior insula (AI) (Deng et al. 2014; Giuliani et al. 2011).

Nonetheless, within the bodies of research on the brain structural correlates both of expertise at skills associated to certain professions, and of the empathic response and its components, no study has sought to compare two groups who—in virtue of their profession—clearly have disparate empathic capacities. For this reason, it is hypothesized that psychotherapists, who are in constant demand of modulating their empathic response, will display brain cortical variations in regions engaged in such cognitive endeavors. Therefore, in this study we compared the cortical thickness at several prefrontal and cingulate regions of a control group to that of a psychotherapists group and determined whether in the latter local cortical thickness was associated with their empathic capacity.

The psychotherapists studied herein were person-centered psychotherapists who incorporate the generation of *alliance* as a fundamental psychotherapeutic tool. Alliance refers to the kind of bonds and synchrony between user and psychotherapist that is achieved through empathic efforts and commitment to the therapeutic process (Horvath 2001; Koole & Tschacher 2016) and which has been consistently associated with therapeutic success (Goldsmith et al. 2015; Klein et al. 2003; McLeod 2011; Webb et al. 2010). Hence, a conscious modulation of the psychotherapist’s empathic response is deemed crucial for the generation of alliance (Evans-Jones et al. 2009; Horvath, 2006). We thus claim that our participants have a greater and more explicit need to constantly engage in effective empathic responses than professionals in other work areas. The relevance of this study lies in providing a further understanding of the adult brain plasticity, by comparing for the first time two populations that differ in their empathic skills and their need to actively build these up.

## Methods

### Participants

Thirty-six participants were recruited for the study, 18 of whom were person-centered psychotherapists (TA; 9 female; age mean 54.9 ±7.6), who were contacted through their affiliation to a local mental health organization. All psychotherapists had graduate studies in clinical psychotherapy as well as at least six years of professional clinical experience and were professionally active at the moment of this study. A semi-structured interview with them was conducted in order to ensure that they explicitly employed *alliance* as a psychotherapeutic tool.

The control group consisted of 18 non-therapists (NT; 9 female; age mean 54.7 ±7.6), who were healthy professionals from the different fields specified in the national classification of occupations. All of them had approximately the same years of studies, socioeconomic status and professional working experience as the psychotherapists group. All participants were right-handed, reported no neurological or psychiatric history, and were not taking any psychotropic medication. Each one signed a written informed consent to take part in the study. This project was approved by the ethics committee of the Neurobiology Institute (UNAM), following the Declaration of Helsinki guidelines.

### Questionnaires

Empathic skills were assessed through the interpersonal reactivity index (IRI; Davis 1983), which is subdivided into four independent dimensions: *Fantasy* (that evaluates the tendency to identify with fictitious characters and situations, e.g. from a book), *Perspective Taking* (measures the tendency to actively adopt another person’s point of view in orders to understand how they feel), *Empathic Concern* (evaluates the tendency to react emotionally to the suffering of the others, with sentiments of compassion, care and concern), and *Personal Distress* (which measures the tendency to experience distress in stressful interpersonal situations, including emergencies) (Carrazco-Ortiz et al. 2011; Davis 1983; Escrivá et al. 2004). Emotion regulation was assessed by the Emotion Regulation Questionnaire (ERQ). Specifically, this questionnaire evaluates the tendency to adopt *Expressive Suppression* and *Cognitive Reappraisal* as emotion modulation strategies (Gross & John 2003; Mauss et al. 2007).

Moreover, all participants completed The Toronto Alexithymia Scale (TAS-20; Bagby et al. 1994), the score of which was used as an exclusion criterion if it suggested pre-clinical or clinical alexithymia. The reason for this is that alexithymic traits have been consistently associated with empathy deficits (Banzhaf et al. 2018; Goerlich et al. 2017).

### MRI acquisition

Structural MR imaging was performed on a 3 Tesla scanner (General Electric, Waukesha, WI), with a 32-channel array head coil. A whole brain three-dimensional high resolution T1 weighted image was acquired from each participant through a spoiled gradient recalled echo sequence (SPGR) with the following parameters: repetition time (TR) = 8.1ms, echo time (TE) = 3.2 ms, flip angle = 12°, isometric voxel = 1.0 × 1.0 × 1.0 mm^3^, image matrix = 256 × 256, FOV = 256, acquisition plane = Sagittal, acceleration factor = 2.

### Preprocessing

A qualitative quality control was performed on each raw MRI volume by trained observers, evaluating four criteria: image sharpness, presence of the ringing artifact, contrast to noise ratio (CNR) on subcortical structures, and CNR around the cortex (Backhausen et al. 2016). In this process, we decided to exclude one volume pertaining to the non-therapists group from further analyses, since it did not display an acceptable quality according to this method. Noise and magnetic field inhomogeneities were corrected using Advanced Normalization Tools (ANTs; Tustison et al. 2010). Finally, a binary mask was generated for each brain image using the online volBrain 1.0 pipeline (Manjón & Coupé 2016). The resulting masks’ quality were also qualitatively controlled, and small manual corrections were applied when needed.

We selected regions of interest (ROIs) relevant to cognitive empathy and emotion regulation from the Human Brainnetome Atlas (BNA; Fan et al. 2016). This was done using the BNA’s Interactive Website Viewer (atlas.brainnetome.org/bnatlas.html), which shows behavioral and paradigm class metadata associated with each region available in the atlas (Fan et al. 2016). Namely, we identified regions associated with the keyword *social cognition* in the prefrontal and cingulate cortices. We then further restricted this selection according to relevant previous literature on cognitive empathy and emotion regulation. Next, the corresponding probabilistic maps were downloaded, thresholded, and binarized to serve as ROI masks. The resulting ROIs are listed in Table 1.

**Table 1.**
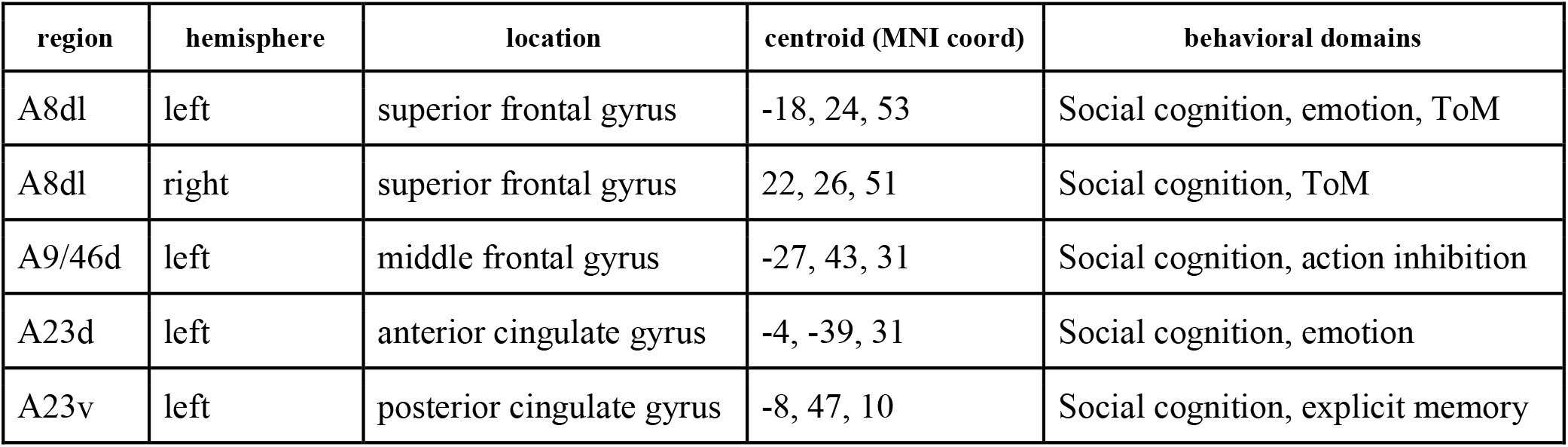
List of ROIs selected from the Human Brainnetome Atlas (Fan et al. 2016) based on their associated behavioral domains and further restricted by previous literature.

| region | hemisphere | location | centroid (MNI coord) | behavioral domains |
| --- | --- | --- | --- | --- |
| A8dl | left | superior frontal gyrus | -18, 24, 53 | Social cognition, emotion, ToM |
| A8dl | right | superior frontal gyrus | 22, 26, 51 | Social cognition, ToM |
| A9/46d | left | middle frontal gyrus | -27, 43, 31 | Social cognition, action inhibition |
| A23d | left | anterior cingulate gyrus | -4, -39, 31 | Social cognition, emotion |
| A23v | left | posterior cingulate gyrus | -8, 47, 10 | Social cognition, explicit memory |

### Cortical thickness measurement

The CIVET 2.1.1 pipeline was used to extract brain surfaces and estimate cortical thickness in each brain image (Montreal Neurological Institute at McGill University, Montreal, Quebec, Canada). A 12-parameter affine transformation was applied to each individual preprocessed image, registering it to the MNI ICBM 152 model. Then a tissue classification was performed, that is, each voxel was assigned to represent white matter (WM), gray matter (GM) or cerebro-spinal fluid (CSF) based on signal intensity and *a priori* probabilistic models. The preprocessed T1-weighted volumes were fed to the pipeline with the individual volBrain masks for better segmentation. Next, deformable models were used to create and extract WM and GM surfaces for each hemisphere separately, yielding 40,952 vertices per surface. Then an estimation of the distances between the internal and the external cortical surfaces were estimated at each vertex, and smoothed using a 25 mm surface-based diffusion blurring kernel. For our study we extracted only cortical thickness data.

### Statistical analysis

We carried out all statistical analyses using R (R Core Team, 2020). Cortical thickness (CT) data were analyzed with the RMINC package (Lerch et al. 2017). Using a linear model, vertex-wise comparisons were performed at each region of interest. Correction for multiple comparisons was made using false discovery rate (FDR; Genovese et al. 2002) at 5%. Age and sex were included in the model as covariates of no interest. Next, another linear model was used to determine whether the asymmetry measures (the difference between corresponding vertices of the two hemispheres) were different between groups. Finally, another linear model was used to explore if cortical thickness at the ROIs varied as a function of any of the psychometric scores. Sex and age were again included as covariates of no interest. The effect size for the first linear model was additionally calculated for the model (in terms of Hedges’ g) at every vertex of the brain surface.

## Results

### Behavioral results

The psychometric results show that psychotherapists tended to score differently than the non-therapists in some of the subscales of the tests applied. The mean score at each IRI and ERQ subscale by group is displayed with its standard deviation in Table 2.

**Table 2.** Mean psychometric scores and their standard deviations by group. Fantasy, Perspective-Taking, Empathic Concern, and Personal Distress belong to the IRI, whereas Cognitive Reappraisal and Expressive Suppression belong are subscales of the ERQ.

| Group | *Fantasy* | *Perspective Taking* | *Empathic Concern* | *Personal Distress* | *Cognitive Reappraisal* | *Expressive Suppression* |
| --- | --- | --- | --- | --- | --- | --- |
| **Therapists** | 19.17 ($\pm 3.75$) | 22.72 ($\pm 3.58$) | 22.61 ($\pm 3.07$) | 8.06 ($\pm 3.33$) | 4.70 ($\pm 1.29$) | 1.89 ($\pm 0.71$) |
| **Non-therapists** | 12.65 ($\pm 5.36$) | 19.12 ($\pm 4.94$) | 21.06 ($\pm 5.56$) | 9.24 ($\pm 5.09$) | 4.33 ($\pm 1.83$) | 2.85 ($\pm 1.29$) |

As can be seen in Figure 1, psychotherapists scored significantly higher than non-therapists on the two IRI subscales related to cognitive empathy: *Fantasy* (p < 0.01) and *Perspective Taking* (p < 0.05). Regarding the other two IRI constructs, no statistically significant difference was found between the groups.

**Fig. 1.**
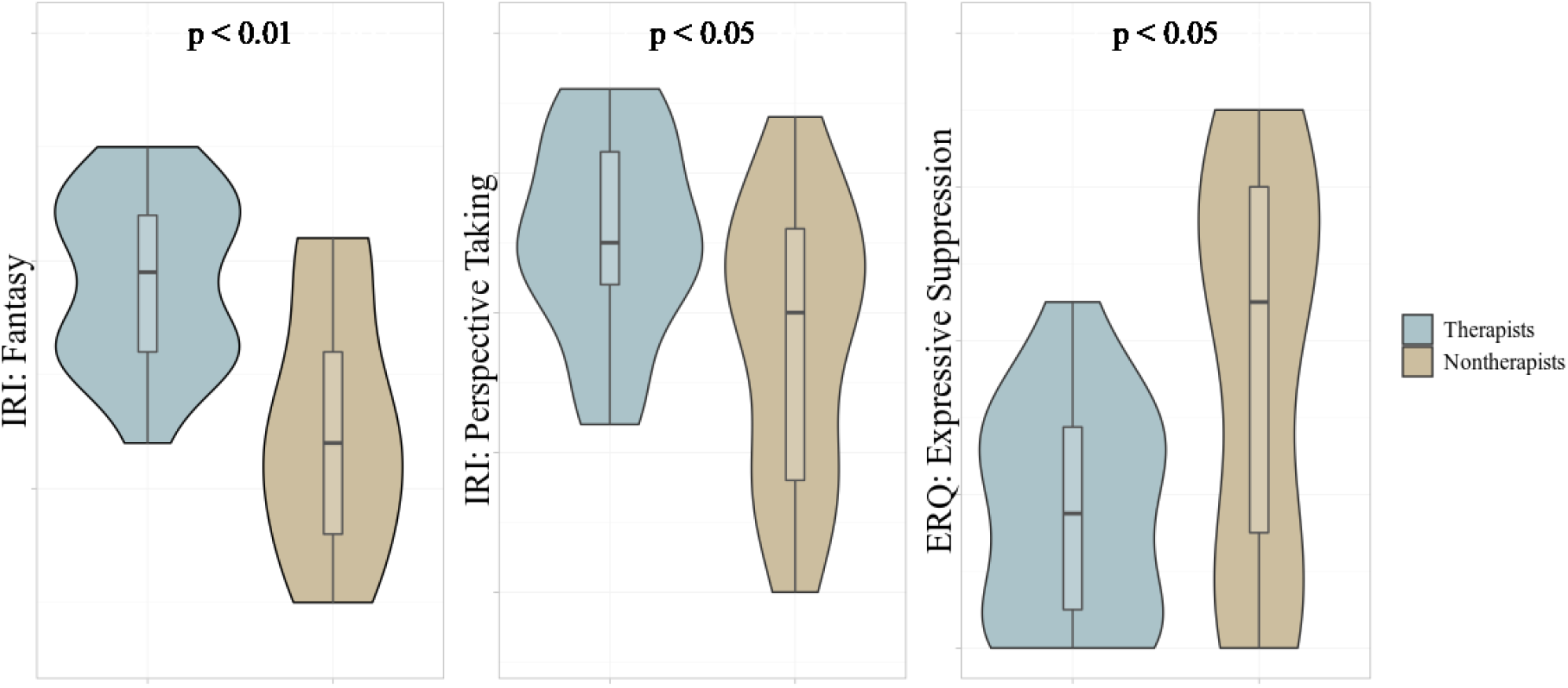
Scores in which statistically significant inter-group differences were found

Psychotherapists also scored significantly lower on the ERQ *Expressive Suppression* construct (p < 0.05) (Figure 1); however, no statistically significant difference was to be found in the *Cognitive Reappraisal* subscale.

### Cortical thickness results

Out of the five ROIs, a statistically significant difference (FDR < 5%) was found in one dorsolateral prefrontal area, displayed in Figure 2. Concretely, the psychotherapists displayed a greater cortical thickness on the left A9/46d region than non-therapists (q < 0.05, FDR corrected). No other significant cortical thickness differences were to be found between the groups in the other prefrontal and cingulate ROIs.

**Fig. 2.**
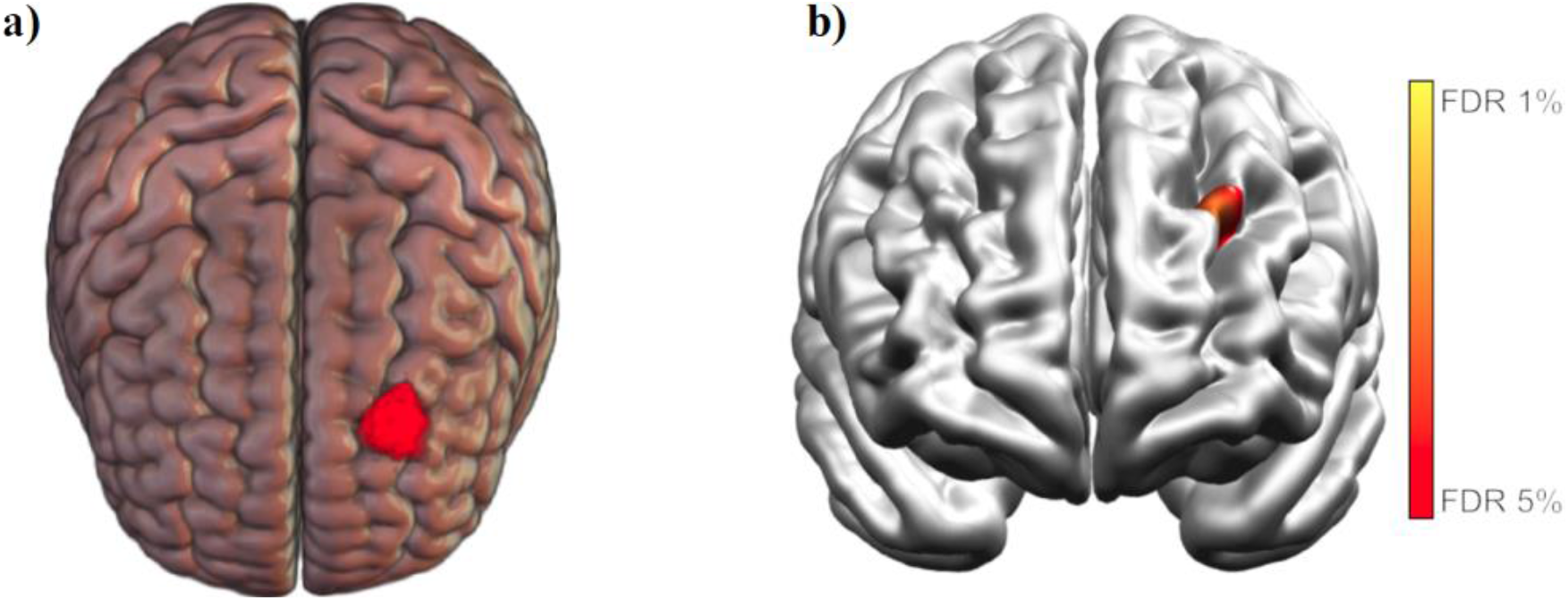
a) left A9/46d region; b) statistical map of left A9/46d region (therapists > non-therapists). (a) was generated with the MRIcroGL software (Rorden & Brett 2000); (b) was generated using the Display software (http://www.bic.mni.mcgill.ca/software/Display/Display.html)

After fitting the vertex-wise linear models for left A9/46d cortical thickness and psychometric scores of the psychotherapists group, we found that their cortical thickness in this region was negatively correlated with their IRI’s *Empathic Concern* score (q < 0.01, FDR corrected). Figure 3 displays a representation of this result; i.e. it presents a linear model of individual mean local cortical thickness and total *Empathic Concern* score. No other statistically significant associations were found between the region’s cortical thickness and psychometric scores.

**Fig. 3.**
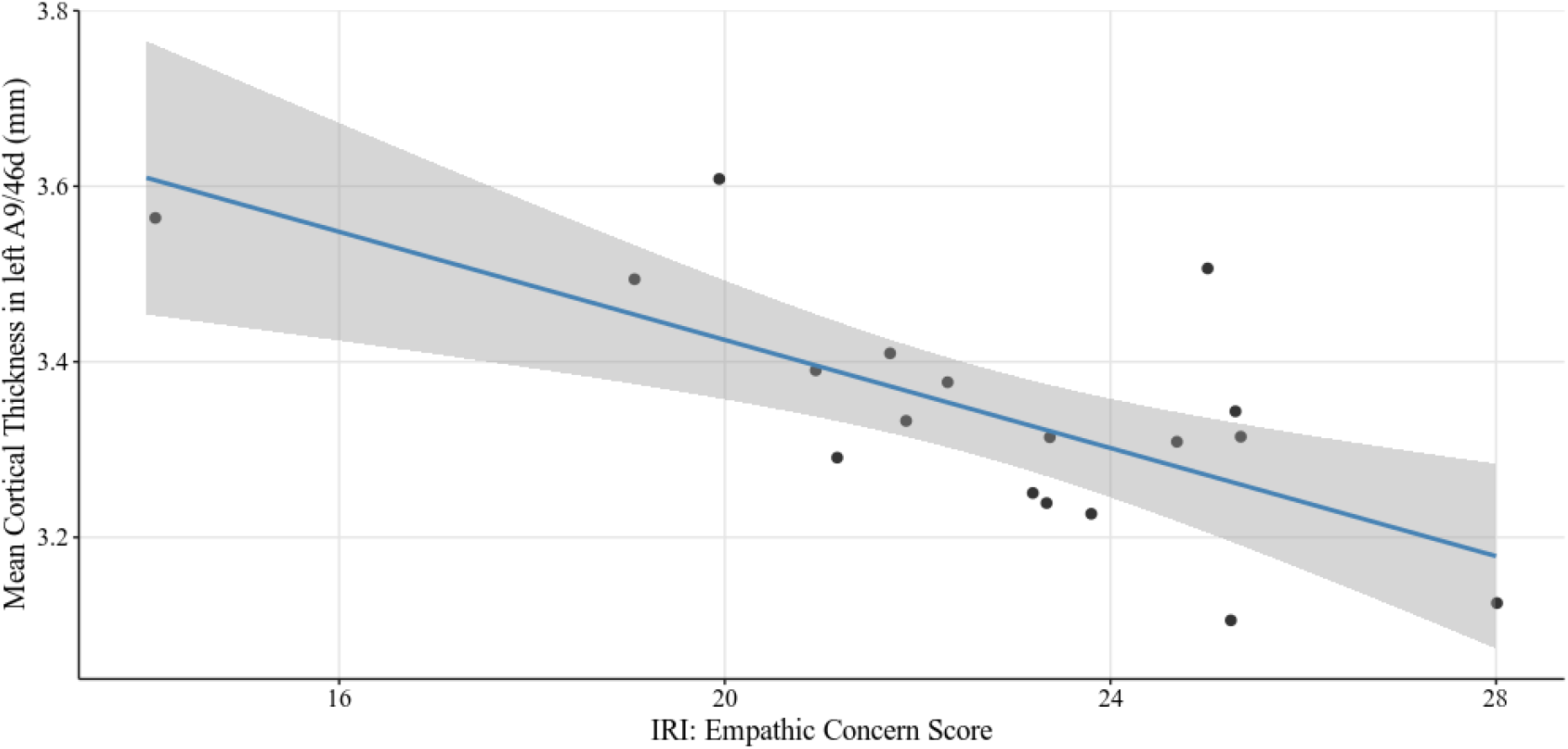
Representation of the significant fitted linear model (individual Empathic Concern score vs. mean left A9/46d mean cortical thickness of the psychotherapists group) with its 95% level confidence interval (in gray)

Furthermore, the vertex-wise cortical thickness asymmetry linear model did also reveal a higher cortical thickness asymmetry at the A9/46d region in the therapists’ brains (left > right) than in the non-therapists’, reaching a higher FDR threshold (q < 0.1, FDR corrected).

Finally, depicted in Figure 4, a whole-brain effect size statistical map was computed for the first (groups comparison) linear model, showing that the same dorsolateral prefrontal cortex region displays the highest effect size values. Although beside the point of this study, it might be worth noting that sensorimotor regions, such as the inferior somatosensory cortex and paracentral gyrus, display a tendency to high effect sizes (g > 8.0).

**Fig. 4.**
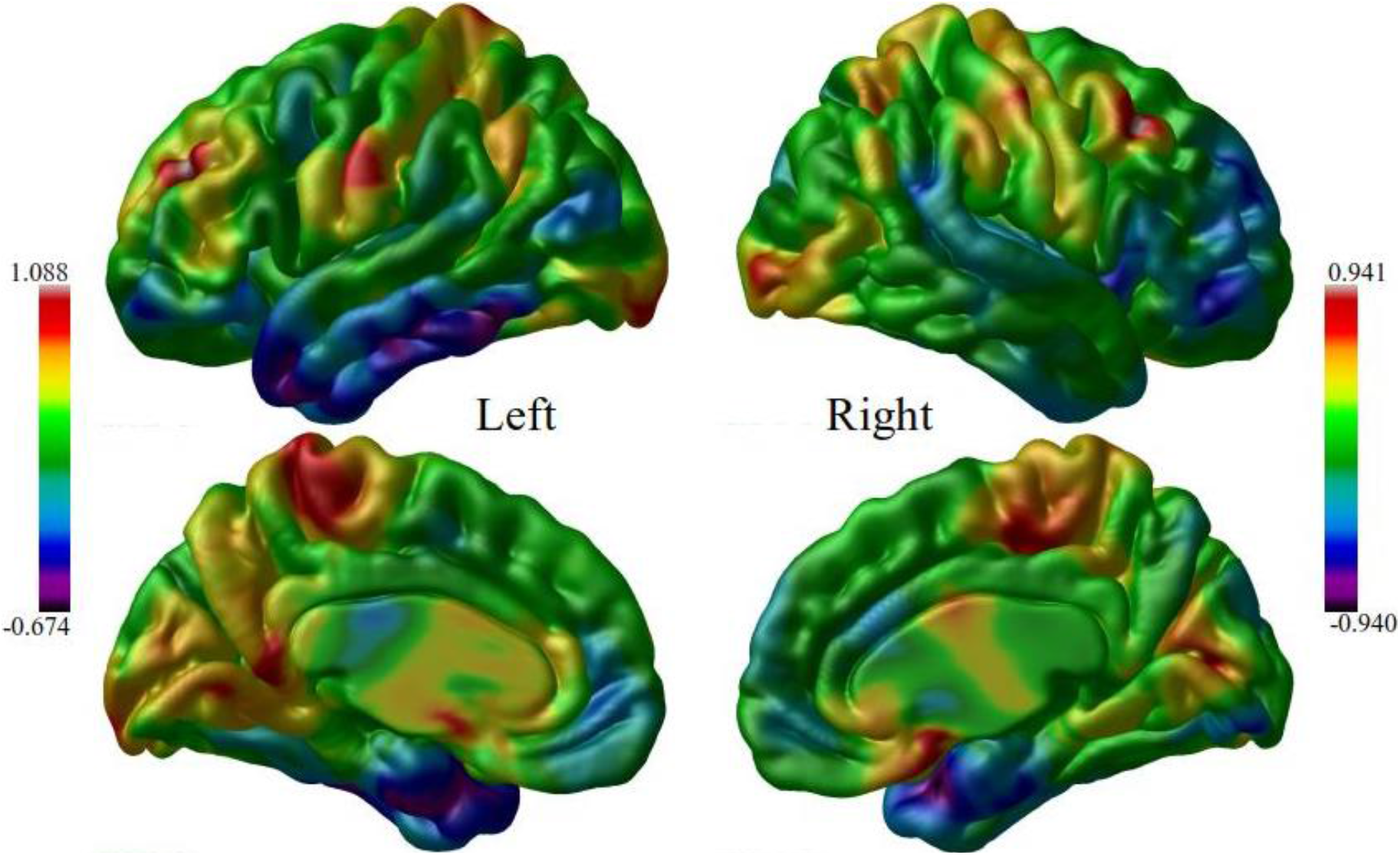
Whole brain effect size statistical map (Hedges’ g) generated from the linear model comparing groups’ cortical thickness measures

## Discussion

The aim of this study was to investigate the cortical thickness variations associated with greater empathic response in a group of psychotherapists. The main finding is that a region of interest in the left dlPFC (A9/46d) was significantly thicker in the group of psychotherapists than in the non-therapists control group. This region, related to executive control, has not only been associated with perspective-taking skills (De Waal & Preston 2017; Völlm et al. 2016), but it has also been deeply implicated in top-down emotion regulation processes (Enzi et al. 2016; Ochsner & Gross 2005). What is more, activation in this region negatively correlates with activity in limbic structures—including the amygdala—during the presentation of emotionally charged stimuli (Sang & Hamman 2007; Smoski et al. 2014; Urry et al. 2006). Therefore, the greater thickness of this region may reflect a greater tendency to regulate one’s affective states.

In addition, we found that psychotherapists’ cortical thickness in this left dlPFC region displays a stronger asymmetry with respect to its homologous in the right hemisphere than the control group. This is consistent with previous literature that has shown that the cognitive components of empathy (i.e. perspective taking) and the empathic response (i.e. emotion regulation) are somewhat left-lateralized (Ochsner et al. 2004).

Another important finding regarding the A9/46d region is that, in psychotherapists, the cortical thickness is negatively correlated with the IRI’s *Empathic Concern* score. The Empathic Concern construct has been repeatedly found to be IRI’s strongest component—i.e. it is the most representative of empathy as a whole (Alterman et al. 2003; Cliffordson 2001). This result suggests that thickness in this cortical region is directly associated with the psychotherapists’ tendency to feel empathically concerned for others and, thus, effectively engage in helping behavior (Lamm, Batson & Decety 2017). In the context of the psychotherapeutic practice, helping behavior could mean consciously modulating one’s own empathic response (Prikhidko & Swank 2018). Moreover, its association with A9/46d is consistent with previous findings which suggest that the tendency to feel empathically concerned for others is correlated with the ability to regulate one’s affective states (Eisenberg, Spinrad & Morris 2013; Lamm et al. 2007b), which is a crucial function of this region. It thus might be that variations in this region could be reflecting the psychotherapists’ tendency to be differently empathic through persistently modulating their affective state.

However, contrary to what would be expected from the dlPFC region, we did not find any significant correlation between the ERQ scores and left A9/46d cortical thickness. This is likely either because the psychotherapists engage in other cognitive emotion regulation mechanisms and strategies not measured here, or because the ERQ could not capture their higher tendency to engage in cognitive reappraisal strategies to regulate their affective states. Either of these alternatives could find support in the fact that they scored significantly lower on the ERQ Expressive Suppression scale with respect to non-therapists. Future research projects may be able to shed light on this issue.

The subsequent, exploratory effect size statistical map revealed that the A9/46d region indeed yields some of the highest effect size values from the model, confirming its prominent role in psychotherapists. On detailed examination, sensorimotor regions also displayed somewhat high effect values; therefore, it could be speculated that these regions might play a role in the neuroplasticity of the psychotherapists’ brain. Since this is in consonance with the perception-action theory of empathy (de Waal & Preston 2017; Preston & de Waal 2002), as well as with previous functional results (Gallo et al. 2018; Engen & Singer 2013), it would be worthwhile for future research efforts to address this issue.

It would also be interesting for future research to measure personality traits, in order to determine whether the detected structural changes are also related to individual differences in personality. Moreover, longitudinal research is required to determine whether psychotherapists develop their increased empathic skills as a result of their professional training and experience, or whether they are drawn to the job because of their already higher empathic capacity. Lastly, this study sample is relatively small due to the difficulty that the recruitment of professional psychotherapists represents. However, given the clearness and robustness of the results, we do not deem that this undermines the study’s value.

## Conclusions

This study indicates that there is a significant difference in cortical thickness at a portion of the dorsolateral prefrontal cortex in a group of psychotherapists as compared to non-therapists, and that psychotherapists’ cortical thickness in this brain region is strongly correlated to their tendency to feel empathically concerned for others. In light of previous literature and of evidence on this region’s functional role, we suggest that these variations reflect individual differences in empathic responding. In conclusion, these results suggest for the first time that a region in the dlPFC presents structural variations associated with empathic capacity in a group of healthy professionals who are required to have a constant active modulation of their empathic response. This adds to the body of studies on neuroanatomical variations associated with different areas of expertise and sheds light on the problem of how the adult brain plasticity responds to environmental demands.

## Declarations

### Ethics approval

This project was approved by the Ethics Committee of the Instituto de Neurobiología, National Autonomous University of Mexico (UNAM). The procedures used in this study adhere to the tenets of the Declaration of Helsinki. Informed consent was obtained from all individual participants included in this study.

## Acknowledgments

We thank Nuri Aranda-López, Leopoldo González-Santos, Ma. de Lourdes Lara Ayala, and Erick H. Pasaye for their technical support, Israel Vaca-Palomares for his valuable advice and support, the UNAM Academic Writing Program team for helping MED develop his writing skills, and M.C. Jeziorski for editing the manuscript. The authors thankfully acknowledge the imaging resources and support provided by the “Laboratorio Nacional de Imagenología por Resonancia Magnética”, CONACyT Network of National Laboratories.

## Data availability

The datasets analyzed during the current study are available from the corresponding author upon request.

## References

Alterman AI, McDermott PA, Cacciola JS & Rutherford MJ (2003) Latent Structure of the Davis Interpersonal Reactivity Index in Methadone Maintenance Patients. Journal of Psychopathology and Behavioral Assessment, 25(4), 257–265. https://doi.org/10.1023/A:1025936213110

Ashburner J Friston KJ (2000) Voxel-based morphometry - The methods. NeuroImage, 11(6 I), 805–821. https://doi.org/10.1006/nimg.2000.0582

Backhausen LL, Herting MM, Buse J, Roessner V, Smolka MN & Vetter NC (2016) Quality control of structural MRI images applied using FreeSurfer-a hands-on workflow to rate motion artifacts. Frontiers in Neuroscience, 10, 1–10. https://doi.org/10.3389/fnins.2016.00558

Bagby M, Parker JD & Taylor GJ (1994) the Twenty-Item Item Selection Toronto and Cross-Validation Structure. Journal of Psychosomatic Research, 38(1), 23–32. https://doi.org/10.1016/0022-3999(94)90005-1

Banissy MJ, Kanai R, Walsh V & Rees G (2012) Inter-individual differences in empathy are reflected in human brain structure. NeuroImage, 62(3), 2034–2039. https://doi.org/10.1016/j.neuroimage.2012.05.081

Banks SJ, Sreenivasan KR, Weintraub DM, Baldock D, Noback M, Pierce ME, Frasnelli J, James J, Beall E, Zhuang X, Cordes D & Leger GC (2016) Structural and functional MRI differences in master sommeliers: A pilot study on expertise in the brain. Frontiers in Human Neuroscience, 10(August), 12. https://doi.org/10.3389/fnhum.2016.00414

Banzhaf C, Hoffmann F, Kanske P, Fan Y, Walter H, Spengler S, Schreiter S, Singer T & Bermpohl F (2018) Interacting and dissociable effects of alexithymia and depression on empathy. Psychiatry Research, 270, 631–638. https://doi.org/10.1016/j.psychres.2018.10.045

Bermudez P & Zatorre RJ (2005) Differences in gray matter between musicians and nonmusicians. Annals of the New York Academy of Sciences, 1060(2005), 395–399. https://doi.org/10.1196/annals.1360.057

Bernhardt BC, & Singer T (2012) The Neural Basis of Empathy. Annual Review of Neuroscience, 35(1), 1–23. https://doi.org/10.1146/annurev-neuro-062111-150536

Carrasco Ortiz MA, Delgado Egido B, Barbero García MI, Holgado Tello FP & del Barrio Gándara MV (2011) Psychometric properties of the Interpersonal Reactivity Index in Spanish child and adolescent population. Psicothema, 23(4), 824–831. http://www.ncbi.nlm.nih.gov/pubmed/22047879

Cliffordson C & Väst H (2001) Parent’s Judgments and Students’ Self-Judgments of Empathy: The Structure of Empathy and Agreement of Judgment Based on the Interpersonal Reactivity Index (IRI) Grades: Comparability, prognostic validity and effects on learning View project Grade and grad. European Journal of Psychological Assessment, December. https://doi.org/10.1027/1015-5759.17.1.36

Davis MH (1983) Measuring individual differences in empathy: Evidence for a multidimensional approach. Journal of Personality and Social Psychology, 44(1), 113–126. https://doi.org/10.1037/0022-3514.44.1.113

de Waal FBM (2012) The Antiquity of Empathy. Science, 336(6083), 874–876. https://doi.org/10.1126/science.1220999

de Waal, FBM (2008) Putting the Altruism Back into Altruism: The Evolution of Empathy. Annual Review of Psychology, 59(1), 279–300. https://doi.org/10.1146/annurev.psych.59.103006.093625

de Waal FBM & Preston SD (2017) Mammalian empathy: Behavioural manifestations and neural basis. Nature Reviews Neuroscience, 18(8), 498–509. https://doi.org/10.1038/nrn.2017.72

Decety J (2011) Dissecting the neural mechanisms mediating empathy. Emotion Review, 3(1), 92–108. https://doi.org/10.1177/1754073910374662

Delon-Martin C, Plailly J, Fonlupt P, Veyrac A & Royet JP (2013) Perfumers’ expertise induces structural reorganization in olfactory brain regions. NeuroImage, 68, 55–62. https://doi.org/10.1016/j.neuroimage.2012.11.044

Deng Z, Wei D, Xue S, Du X, Hitchman G & Qiu J (2014) Regional gray matter density associated with emotional conflict resolution: Evidence from voxel-based morphometry. Neuroscience, 275, 500–507. https://doi.org/10.1016/j.neuroscience.2014.06.040

Eisenberg N, Spinrad TL & Morris A (2013) Empathy-Related Responding in Children. In M. Killen & J. G. Smetana (Eds.), Handbook of Moral Development (2nd ed., pp. 184–207). Psychology Press. https://doi.org/10.4324/9780203581957

Elliott R, Bohart AC, Watson JC & Greenberg LS (2011). Empathy. Psychotherapy, 48(1), 43–49. https://doi.org/10.1037/a0022187

Engen HG & Singer T (2013) Empathy circuits. Current Opinion in Neurobiology, 23(2), 275–282. https://doi.org/10.1016/j.conb.2012.11.003

Enzi B, Amirie S & Brüne M (2016) Empathy for pain-related dorsolateral prefrontal activity is modulated by angry face perception. Experimental Brain Research, 234(11), 3335–3345. https://doi.org/10.1007/s00221-016-4731-4

Eres R, Decety J, Louis WR & Molenberghs P (2015) Individual differences in local gray matter density are associated with differences in affective and cognitive empathy. NeuroImage, 117, 305–310. https://doi.org/10.1016/j.neuroimage.2015.05.038

Escrivá VM, Navarro MDF & García PS (2004) Measuring empathy: The interpersonal reactivity index. Psicothema, 16(2), 255–260.

Etkin A, Büchel C & Gross JJ (2015) The neural bases of emotion regulation. Nature Reviews Neuroscience, 16(11), 693–700. https://doi.org/10.1038/nrn4044

Evans-Jones C, Peters E, & Barker C (2009) The therapeutic relationship in CBT for psychosis: Client, therapist and therapy factors. Behavioural and Cognitive Psychotherapy, 37(5), 527–540. https://doi.org/10.1017/S1352465809990269

Fan L, Li H, Zhuo J, Zhang Y, Wang J, Chen L, Yang Z, Chu C, Xie S, Laird AR, Fox PT, Eickhoff SB, Yu C & Jiang T (2016) The Human Brainnetome Atlas: A New Brain Atlas Based on Connectional Architecture. Cerebral Cortex, 26(8), 3508–3526. https://doi.org/10.1093/cercor/bhw157

Gallo S, Paracampo R, Müller-Pinzler L, Severo MC, Blömer L, Fernandes-Henriques C, Henschel, A, Lammes BK, Maskaljunas T, Suttrup J, Avenanti A, Keysers C & Gazzola, V (2018) The causal role of the somatosensory cortex in prosocial behaviour. ELife, 7, 1–31. https://doi.org/10.7554/eLife.32740

Genovese CR, Lazar NA & Nichols T (2002). Thresholding of statistical maps in functional neuroimaging using the false discovery rate. NeuroImage 15 (4), 870–878

Giuliani NR, Drabant EM, Bhatnagar R & Gross JJ (2011) Emotion regulation and brain plasticity: Expressive suppression use predicts anterior insula volume. NeuroImage, 58(1), 10–15. https://doi.org/10.1016/j.neuroimage.2011.06.028

Goerlich KS, Votinov M, Lammertz SE, Winkler L, Spreckelmeyer KN, Habel U, Gründer G & Gossen A (2017) Effects of alexithymia and empathy on the neural processing of social and monetary rewards. Brain Structure and Function, 222(5), 2235–2250. https://doi.org/10.1007/s00429-016-1339-1

Goldsmith LP, Lewis SW, Dunn G, & Bentall RP (2015). Psychological treatments for early psychosis can be beneficial or harmful, depending on the therapeutic alliance: An instrumental variable analysis. Psychological Medicine, 45(11), 2365–2373. https://doi.org/10.1017/S003329171500032X

Gross JJ (2002) Emotion regulation: Affective, cognitive, and social consequences. Psychophysiology, 39(3), 281–291. https://doi.org/10.1017/S0048577201393198

Hassenstab J, Dziobek I, Rogers K, Wolf OT & Convit A (2007) Knowing what others know, feeling what others feel: A controlled study of empathy in psychotherapists. Journal of Nervous and Mental Disease, 195(4), 277–281. https://doi.org/10.1097/01.nmd.0000253794.74540.2d

Horvath AO (2001) The Alliance. Psychotherapy Theory Research & Practice, 38(4), 365–372. https://doi.org/10.1037/0033-3204.38.4.365

Horvath AO (2006) The alliance in context: Accomplishments, challenges, and future directions. Psychotherapy, 43(3), 258–263. https://doi.org/10.1037/0033-3204.43.3.258

Klein DN, Schwartz JE, Santiago NJ, Vivian D, Vocisano C, Arnow B, Manber R, Riso LP, McCullough JP, Borian FE, Castonguay LG, Blalock JA, Markowitz JC, Rothbaum B, Thase ME, Miller IW & Keller MB (2003) Therapeutic Alliance in Depression Treatment: Controlling for Prior Change and Patient Characteristics. Journal of Consulting and Clinical Psychology, 71(6), 997–1006. https://doi.org/10.1037/0022-006X.71.6.997

Koole SL & Tschacher W (2016) Synchrony in psychotherapy: A review and an integrative framework for the therapeutic alliance. Frontiers in Psychology, 7(June), 1–17. https://doi.org/10.3389/fpsyg.2016.00862

Lamm C, Batson CD & Decety J (2007a) The neural substrate of human empathy: Effects of perspective-taking and cognitive appraisal. Journal of Cognitive Neuroscience, 19(1), 42–58. https://doi.org/10.1162/jocn.2007.19.1.42

Lamm C, Nausbaum HC, Meltzoff AN & Decety J (2007b) What are you feeling? Using functional magnetic resonance imaging to assess the modulation of sensory and affective responses during empathy for pain. PLoS ONE, 2(12). https://doi.org/10.1371/journal.pone.0001292

Lerch JP & Evans AC (2005) Cortical thickness analysis examined through power analysis and a population simulation. NeuroImage, 24(1), 163–173. https://doi.org/10.1016/j.neuroimage.2004.07.045

Lerch J, Hammill C, van Eede M & Cassel D (2017). RMINC: Statistical Tools for Medical Imaging NetCDF (MINC) Files. R package version 1.5.2.1, http://mouse-imaging-centre.github.io/RMINC.

Lerch JP, Van Der Kouwe AJW, Raznahan A, Paus T, Johansen-Berg H, Miller KL, Smith SM, Fischl B & Sotiropoulos SN (2017) Studying neuroanatomy using MRI. Nature Neuroscience, 20(3), 314–326. https://doi.org/10.1038/nn.4501

MacFarlane P, Anderson T & McClintock AS (2017) Empathy from the client’s perspective: A grounded theory analysis. Psychotherapy Research, 27(2), 227–238. https://doi.org/10.1080/10503307.2015.1090038

Maguire EA Woollett K & Spiers HJ (2006) London taxi drivers and bus drivers: A structural MRI and neuropsychological analysis. Hippocampus, 16(12), 1091–1101. https://doi.org/10.1002/hipo.20233

Manjón JV & Coupé P (2016) Volbrain: An online MRI brain volumetry system. Frontiers in Neuroinformatics, 10(JUL), 1–14. https://doi.org/10.3389/fninf.2016.00030

McLeod BD (2011) Relation of the alliance with outcomes in youth psychotherapy: A meta-analysis. Clinical Psychology Review, 31(4), 603–616. https://doi.org/10.1016/j.cpr.2011.02.001

Ochsner KN & Gross JJ (2005) The cognitive control of emotion. Trends in Cognitive Sciences, 9(5), 242–249. https://doi.org/10.1016/j.tics.2005.03.010

Ochsner KN, Knierim K, Ludlow DH, Hanelin J, Ramachandran T, Glover G & Mackey SC (2004) Reflecting upon feelings: An fMRI study of neural systems supporting the attribution of emotion to self and other. Journal of Cognitive Neuroscience, 16(10), 1746–1772. https://doi.org/10.1162/0898929042947829

Olalde-Mathieu V, Sassi F, Reyes-Aguilar A, Mercadillo R, Alcauter S & Barrios F (2020) Greater Empathic Abilities and Their Correlation With Resting State Brain Connectivity in Psychotherapists Compared To Non-Psychotherapists. https://doi.org/10.1101/2020.07.01.182998

Pletzer JL, Sanchez X & Scheibe S (2015) Practicing psychotherapists are more skilled at downregulating negative emotions than other professionals. Psychotherapy, 52(3), 346–350. https://doi.org/10.1037/a0039078

Preston SD (2013) The origins of altruism in offspring care. Psychological Bulletin, 139(6), 1305–1341. https://doi.org/10.1037/a0031755

Preston, SD & de Waal FBM (2002) Empathy: Its ultimate and proximate bases. Behavioral and Brain Sciences, 25(1), 1–20. https://doi.org/10.1017/S0140525X02000018

Preston SD & De Waal FBM (2017) Only the PAM explains the personalized nature of empathy. Nature Reviews Neuroscience, 18(12), 769. https://doi.org/10.1038/nrn.2017.140

Prikhidko A & Swank JM (2018) Emotion Regulation for Counselors. Journal of Counseling and Development, 96(2), 206–212. https://doi.org/10.1002/jcad.12193

R Core Team (2020) R: A language and environment for statistical computing. R Foundation for Statistical Computing, Vienna, Austria. URL https://www.R-project.org/

Rorden C & Brett M (2000) Stereotaxic display of brain lesions. Behavioural Neurology, 12, 191–200. www.fil.ion.ucl.ac.uk/spm/

Sang HK & Hamann S (2007) Neural correlates of positive and negative emotion regulation. Journal of Cognitive Neuroscience, 19(5), 776–798. https://doi.org/10.1162/jocn.2007.19.5.776

Smoski MJ, Keng SL, Ji JL, Moore T, Minkel J & Dichter GS (2014) Neural indicators of emotion regulation via acceptance vs reappraisal in remitted major depressive disorder. Social Cognitive and Affective Neuroscience, 10(9), 1187–1194. https://doi.org/10.1093/scan/nsv003

Tustison, NJ, Avants BB, Cook PA, Yuanjie Zheng EA, Yushkevich PA & Gee JC (2010) N4ITK: Improved N3 Bias Correction. IEEE Transactions on Medical Imaging, 29(6), 1310–1320. https://doi.org/10.1109/TMI.2010.2046908

Uribe C, Puig-Davi A, Abos A, Baggio HC, Junque C & Segura B (2019) Neuroanatomical and functional correlates of cognitive and affective empathy in young healthy adults. Frontiers in Behavioral Neuroscience, 13(May), 1–8. https://doi.org/10.3389/fnbeh.2019.00085

Urry HL, Van Reekum CM, Johnstone T, Kalin NH, Thurow ME, Schaefer HS, Jackson CA, Frye CJ, Greischar LL, Alexander AL & Davidson RJ (2006) Amygdala and ventromedial prefrontal cortex are inversely coupled during regulation of negative affect and predict the diurnal pattern of cortisol secretion among older adults. Journal of Neuroscience, 26(16), 4415–4425. https://doi.org/10.1523/JNEUROSCI.3215-05.2006

Völlm BA, Taylor ANW, Richardson P, Corcoran R, Stirling J, McKie S, Deakin JFW & Elliott R (2006) Neuronal correlates of theory of mind and empathy: A functional magnetic resonance imaging study in a nonverbal task. NeuroImage, 29(1), 90–98. https://doi.org/10.1016/j.neuroimage.2005.07.022

Webb CA, DeRubeis RJ & Barber JP (2010) Therapist Adherence/Competence and Treatment Outcome: A Meta-Analytic Review. Journal of Consulting and Clinical Psychology, 78(2), 200–211. https://doi.org/10.1037/a0018912

Zaki J, Weber J, Bolger N y Ochsner K (2009) The neural bases of empathic accuracy. Proceedings of the National Academy of Sciences of the United States of America, 106(27), 11382–11387. https://doi.org/10.1073/pnas.0902666106

Zaki J & Ochsner K (2012) The neuroscience of empathy: Progress, pitfalls and promise. Nature Neuroscience, 15(5), 675–680. https://doi.org/10.1038/nn.3085

